# Machine learning approaches to predict the plant-associated phenotype of *Xanthomonas* strains

**DOI:** 10.1101/2021.05.25.445602

**Authors:** Dennie te Molder, Wasin Poncheewin, Peter J. Schaap, Jasper J. Koehorst

## Abstract

The genus *Xanthomonas* has long been considered to consist predominantly of plant pathogens, but over the last decade there has been an increasing number of reports on non-pathogenic and endophytic members. As *Xanthomonas* species are prevalent pathogens on a wide variety of important crops around the world, there is a need to distinguish between these plant-associated phenotypes. To date a large number of *Xanthomonas* genomes have been sequenced, which enables the application of machine learning (ML) approaches on the genome content to predict this phenotype. Until now such approaches to the pathogenomics of *Xanthomonas* strains have been hampered by the fragmentation of information regarding strain pathogenicity over many studies. Unification of this information into a single resource was therefore considered to be an essential step. Mining of 39 papers considering both plant-associated phenotypes, allowed for a phenotypic classification of 578 *Xanthomonas* strains. For 65 plant-pathogenic and 53 non-pathogenic strains the corresponding genomes were available and *de novo* annotated for the presence of Pfam protein domains used as features to train and compare three ML classification algorithms; CART, Lasso and Random Forest. Recursive feature extraction provided further insights into the virulence enabling factors, but also yielded domains linked to traits not present in pathogenic strains.

## Introduction

The genus of *Xanthomonas* is mostly known for its pathogenic behaviour, with significant economic and agricultural impact (1). *Xanthomonas* spp. infect a wide variety of plant crops, see Table 1 for examples, however individual *Xanthomonas* pathovars usually show a high degree of both host and tissue specificity (2). Whilst non-pathogenic xanthomonads have been reported as early as 1985 (3), during the last decade many new non-pathogenic strains have been discovered (4–9) and it has become apparent that these non-pathogenic strains form an integral part of the *Xanthomonas* epidemic population structure (10). Moreover, non-pathogenic strains show, in comparison to their pathogenic counterparts, a higher level of genetic diversity (1) suggesting that non-pathogenic xanthomonads are generalists that can epiphytically survive on a much wider host range and might play important roles in the microbiome of healthy, asymptomatic hosts (11, 12). While the relative abundance of these non-pathogenic strains is still not known, their undisputable existence has raised the concern that diagnostic misidentifications might result in unnecessary control measures and/or high economic losses (8). The is-not-pathogenic label depends heavily on the tested conditions used. For example, a set of *X. arboricola* pv. fragariae strains isolated from infected strawberries did, when sprayed onto new plants, not cause symptoms (13), but a repetition of the same assay at increased humidity conditions did reveal the pathogenicity of these strains (14). Examples like this underline the importance of testing strains on a large range of hosts and conditions. However, as current tests are all aimed at establishing pathogenicity and given the large number of environmental parameters that impact infection, it is important to integrate extensive testing of strains with a pathogenomics framework providing insight in the relative importance of genome encoded virulence factors.

**Table 1.** Overview of the *Xanthomonas* phenotype database.

| Species | Strains | + | - | Top 5 Most Tested Hosts | Reference |
| --- | --- | --- | --- | --- | --- |
| *X. arboricola* | 258 | 180 | 232 | *Juglans regia*, *Prunus persica*, *Phaseolus vulgaris*,<br>*Fragaria ananassa*, *Capsicum annuum* | (6, 8, 10, 13, 14, 21, 27–39) |
| *X. oryzae* | 70 | 68 | 2 | *Oryza sativa* | (5, 9) |
| *X. translucens* | 58 | 72 | 14 | *Lolium multiflorum*, *Asparagus virgatus*,<br>*Hordeum vulgare*, *Anthurium andreaeanum* | (38, 40–42) |
| *X. campestris* | 40 | 45 | 22 | *Brassica oleracea*, *Musa acuminata*,<br>*Saccharum officinarum*, *Zea mays*, *Prunus persica* | (6, 9, 35, 36, 38, 43, 44) |
| *X. albilineans* | 27 | 18 | 9 | *Saccharum officinarum*, *Zea mays* | (45–47) |
| *X. axonopodis* | 20 | 12 | 14 | *Manihot esculenta*, *Musa acuminata*,<br>*Saccharum officinarum*, *Zea mays*, *Lycopersicon esculentum* | (28, 35, 38, 48) |
| *X. dyei* | 13 | 12 | 27 | *Dysoxylum spectabile*, *Laurelia novae-zelandiae*,<br>*Metrosideros excelsa* | (39) |
| *X. vasicola* | 6 | 12 | 6 | *Musa acuminata*, *Saccharum officinarum*, *Zea mays* | (35) |
| *X. cannabidis* | 5 | 5 | 2 | *Phaseolus vulgaris*, *Capsicum annuum*, *Hordeum vulgare* | (22, 24, 44) |
| *X. sontii* | 5 | 2 | 3 | *Oryza sativa* | (5, 49) |
| *X. floridensis* | 4 | 0 | 8 | *Brassica oleracea*, *Nasturtium officinale* | (50) |
| *X. maliensis* | 4 | 0 | 4 | *Oryza sativa* | (9, 23) |
| *X. fragariae* | 3 | 3 | 0 | *Fragaria ananassa* | (13) |
| *X. euvesicatoria* | 2 | 2 | 0 | *Lycopersicon esculentum*, *Capsicum annuum* | (6) |
| *X. hortorum* | 2 | 0 | 2 | *Daucus carota* | (38) |
| *X. nasturtii* | 2 | 2 | 2 | *Brassica oleracea*, *Nasturtium officinale* | (50) |
| *X. pseudoalbilineans* | 2 | 1 | 1 | *Saccharum officinarum* | (46, 51) |
| *X. sacchari* | 2 | 0 | 2 | *Citrus sinensis*, *Oryza sativa* | (7, 38) |
| *X. hyacinthi* | 1 | 2 | 0 | *Hyacinthus orientalis*, *Scilla tubergeniana* | (52) |
| *X. melonis* | 1 | 0 | 1 | *Citrus sinensis* | (38) |
| *X. theicola* | 1 | 1 | 0 | *Camellia sinensis* | (53) |
| *X. spp.* | 52 | 66 | 41 | *Asparagus virgatus*, *Hordeum vulgare*, *Oryza sativa*,<br>*Nicotiana tabacum*, *Phaseolus vulgaris* | (9, 13, 35, 38, 42) |
| **Total** | 578 | 503 | 392 | 77 Distinct host species | 39 Unique references |

Strains: number of unique strains assayed; +: Number of positive tests; -: number of negative tests.

A vast array of genomic factors have already been found to impact virulence (reviewed in (15, 16)). Many are located in so called pathogenicity clusters such as the *hrp*-cluster expressing type III secretion systems (T3SS) and associated effectors (T3E) (17), the *xps*-cluster coding for a type II secretion system for secretion of host cell wall degrading enzymes (18), the *gum*-cluster responsible for production of the xanthan-based biofilm unique for *Xanthomonas* spp. (19) and the regulation of pathogenicity factors or *rpf*-cluster which positively regulates virulence (20). Elements of many of these clusters are also present in the genomes of the non-pathogens and currently it remains unclear what combinations of features drive the switch in life-style (15). Studies examining the genomic differences between pathogenic and non-pathogenic xanthomonads have focused on *X. arboricola* as a model (21) and as a result the majority of known non-pathogenic strains currently belong to this species. Using classical comparative genomic approaches, attempts have been made to understand what exactly separates pathogenic and non-pathogenic *X. arboricola* strains. These analyses revealed genome encoded differences in environmental sensing, flagellin protein sequences, and components of the type IV pilus, but the most remarkable differences were found in the T3SS and T3E genes content. Most non-pathogenic strains lacked parts of the T3SS and showed a more limited repertoire of T3Es and in extreme cases, non-pathogenic strains even lacked the entire T3SS (1, 8, 11, 22). However, these findings do not fully explain the differences between the two plant associated phenotypes in *X. arboricola*, as strains CFBP3122 and CFBP3123 were found to be pathogenic although they lacked T3SS and T3E genes (11). These findings also provide an incomplete framework for other species. For example, whilst for *X. maliensis* absence of the T3SS related genes appear to be strongly correlated with the non-pathogenic phenotype (23), several members of the *X. cannabis* species have been shown to be pathogenic to multiple different hosts even though they lack the entire T3SS.

Overall the results suggest that a more complete set of genome-encoded features is required to differentiate between phytopathogenic and non-pathogenic strains (22, 24). In the last ten years, the number of available *Xanthomonas* genomes has increased nearly 100-fold (2, 25). This large number of genomes makes it feasible to use machine learning (ML) approaches on the genome content to predict the pathogenicity of individual strains. Until now such approaches have been hampered by the fragmentation of information on the plant-associated phenotype of individual *Xanthomonas* strains over many literature sources. There are databases that track the pathogenicity of individual strains such as CIRM-CFBP (26), but they suffer from poor interoperability and a lack of provenance. Unification of plant-associated phenotype data into a single high quality resource was therefore considered an essential first step.

In this study, pathogenicity assays were retrieved from 39 recent studies that each took into account both pathogenic and non-pathogenic xanthomonads and unified into a single database. This database was then leveraged to extract available genome sequence which were *de novo* annotated for the presence of Pfam protein domains. These domains were subsequently used as feature input to train three different classifiers. The resulting models were examined for their ability to predict the pathogenicity of an individual strain. Important input features were extracted from these models providing new insights into the genome encoded factors contributing to the plant-associated phenotype. At the same time, these classifiers provide a way to cross-validate pathogenicity test results and conditions.

## Results

The workflow is presented in Figure 1. Briefly, pathogenicity assays were obtained from literature and stored in an SQL database. From these assays strains were divided in two classes, available genomes were retrieved and de novo annotated for Pfam protein domains. Domain annotations were used as input to create a strain domain matrix as feature input to train three different classifiers: Classification And Regression Tree (CART), Lasso and Random Forest (RF).

**Fig. 1.**
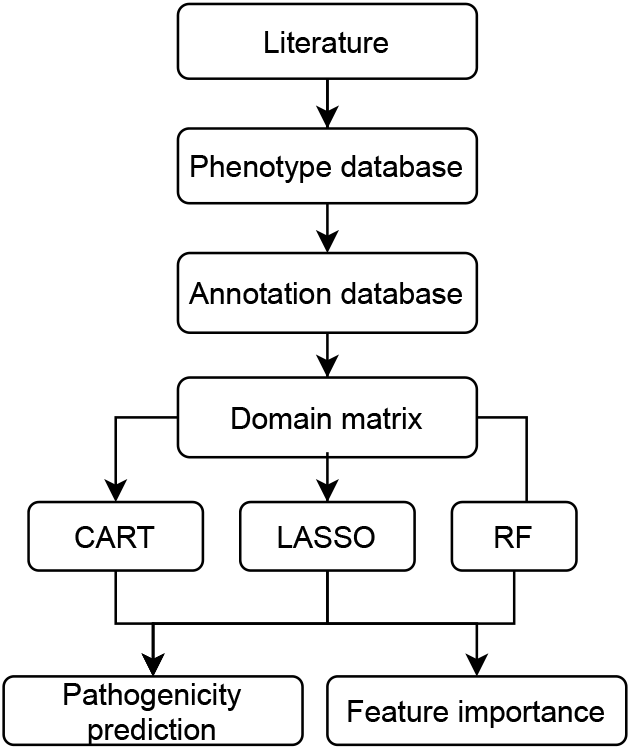
Workflow. *Xanthomonas* pathogenicity assay data and strains were obtained from literature and stored in a SQL (phenotype) database. Available genomes were retrieved and *de novo* annotated for protein domains. Annotation results were stored in a Graph database. Strain specific domain content was used as input to train the classifiers. Resulting models were examined for their ability to predict pathogenicity and feature importance.

### Development of the *Xanthomonas* Phenotype Database

To manage literature derived information related to the plant-associated phenotype, an SQL database was created (Figure S1). This database, which we will be referred to as the phenotype database, was used to track the outcome of individual pathogenicity assays. An assay was defined as the unique combination of a strain and host species as tested by a single source. This approach was favoured over tracking the pathogenicity of individual strains, as it enabled us to also track the criteria used to determine the plant-associated phenotype of a strain. To balance the phenotype database, the data collection effort was limited to recent studies that took both plant-associated phenotypes and their sources into account. This yielded a total of 895 distinct pathogenicity assays, extracted from 39 studies, describing 578 unique strains that were tested on 77 different plant host species (Table 1). Out of the collected 895 individual assays, 503 did and 392 did not indicate strain pathogenicity. From the 578 unique strains, 522 were assayed on their host of isolation (Supplementary File S1).

SQL queries were used to combine results from the various pathogenicity assays. From these combined results the plant-associated phenotype of each individual strain was inferred. Strains unable to induce symptoms on the isolation host after artificial inoculation under optimal conditions were labelled as non-pathogenic (10). Conversely, strains were considered pathogenic if they were able to induce symptoms on any of the tested hosts. Pathogenicity assays based on the “trunk incision” method (31) and pathogenicity assays on *Fragaria ananassa* were both considered to provide insufficient prove for non-pathogenicity, as the former is known to misclassify pathogenic strains that can only cause vertical oozing canker (6, 28) and for the latter the concern was raised that the host might be an unsuitable host to determine pathogenicity of *X. arboricola* strains (13).

Using these criteria, 158 strains were classified as non-pathogenic and 391 strains as pathogenic. Strain names and known aliases of these strains were cross-referenced with the GenBank sequence database (25), to obtain available matching genomes. This resulted in a set 65 pathogenic and 53 non-pathogenic *Xanthomonas* strains with a known genome sequence encompassing a large majority of the observed genetic variation within this genus, with the exception of the of the *X. hortorum, X. gardneri, X. citri, X. perforans* and *X. vesicatoria* species (Table 2).

**Table 2.** Sequenced *Xanthomonads* with a known phenoype.

| Species | Non-Pathogenic | Pathogenic | Total |
| --- | --- | --- | --- |
| *X. arboricola* | 28 | 26 | 54 |
| *X. campestris* | 2 | 6 | 8 |
| *X. oryzae* | 0 | 7 | 7 |
| *X. translucens* | 0 | 7 | 7 |
| *X. cannabidis* | 2 | 3 | 5 |
| *X. vasicola* | 0 | 5 | 5 |
| *X. sontii* | 3 | 1 | 4 |
| *X. albilineans* | 2 | 2 | 4 |
| *X. axonopodis* | 2 | 2 | 4 |
| *X. sacchari* | 2 | 0 | 2 |
| *X. pseudoalbilineans* | 1 | 1 | 2 |
| *X. fragariae* | 0 | 2 | 2 |
| *X. dyei* | 1 | 0 | 1 |
| *X. floridensis* | 1 | 0 | 1 |
| *X. maliensis* | 1 | 0 | 1 |
| *X. melonis* | 1 | 0 | 1 |
| *X. hyacinthi* | 0 | 1 | 1 |
| *X. nasturtii* | 0 | 1 | 1 |
| *X. theicola* | 0 | 1 | 1 |
| *X. spp.* | 7 | 0 | 7 |
| **Total** | 53 | 65 | 118 |

### *De novo* Genome Annotation

Collected genomes were *de novo* annotated for Pfam domains using the SAPP platform (54), which implements Prodigal for gene calling (55) and InterProScan for domain annotation (56). This was done to rule out technical differences due to the use of different annotation software or different versions of the same software. Genome annotations and provenance were stored in a separate database. The statistics are summarised in Table 3. More than 80% of the protein encoding genes code for at least one Pfam domain. The maximum number of genes is inflated by two outlier genomes of low assembly quality, resulting in the prediction of small incomplete genes. However these small genes did not code for protein domains. For annotation details see (Figure S2 and Supplementary File S2).

**Table 3.**
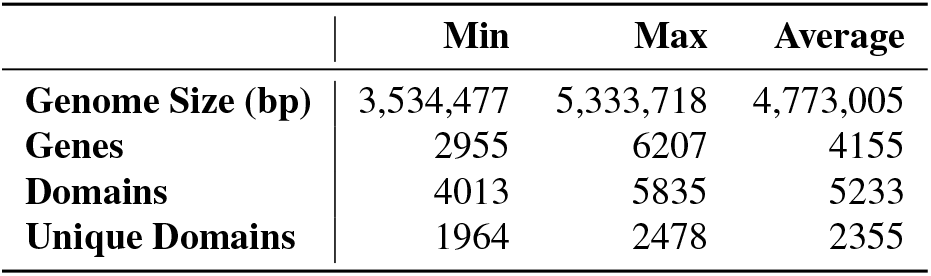
Annotation statistics of the collected genomes.

|  | Min | Max | Average |
| --- | --- | --- | --- |
| **Genome Size (bp)** | 3,534,477 | 5,333,718 | 4,773,005 |
| **Genes** | 2955 | 6207 | 4155 |
| **Domains** | 4013 | 5835 | 5233 |
| **Unique Domains** | 1964 | 2478 | 2355 |

### Domain Matrix

For each strain the set of unique protein domains were extracted from the annotation database and combined into binary strain/domain presence matrix which was used for further analysis and model building. The resulting matrix contained 3609 unique Pfam protein domains distributed over the 118 *Xanthomonas* strains

As the ML models will be developed from the present *Xanthomonas* matrix, a “closed” domain representation of genus is a prerequisite for reliable model building. When a genus is closed, it is assumed that majority of the observed genetic variation within is captured meaning that when new genomes are added, only limited number of new domains will be discovered. The Heaps law estimate was used to estimate the increase in the number of unique domains as a function of the increase in collection size. The decay parameter, alpha, was estimated to be 1.22 indicating that the pan-domainome captured in the matrix was closed (Figure 2). The captured phylogenetic diversity was additionally visualised using a binary domain distance tree (Figure S3 and Supplementary File S2).

**Fig. 2.**
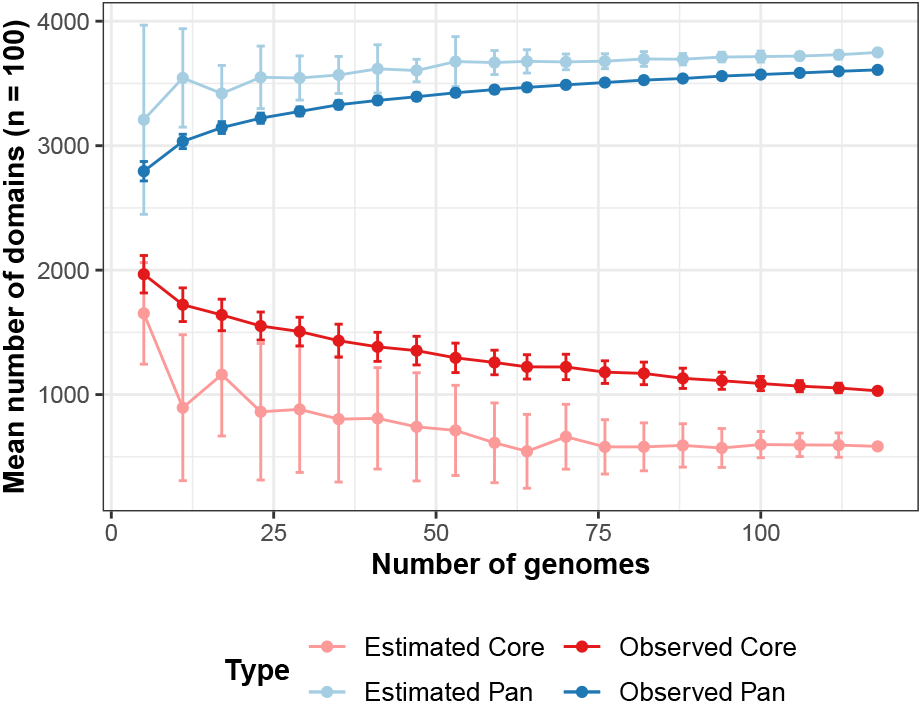
The *Xanthomonas* Pan-domainome is closed. The decay parameter, alpha, was estimated to be 1.22 (see methods for details).

#### Matrix optimization

To reduce the complexity of the data set, the domain matrix was filtered for domains with a low level of variability (present /absent in > 97.5% of samples). These domains either are part of the functional core and thus present in all strains (Figure 2) or represent rare domains containing little information about the general tendencies that discriminate pathogens from non-pathogens. Removal of these domains reduced the total number of domains to 1692. Next, highly correlated domains were treated as one by removal of highly correlated domains with a pair-wise Pearson correlation >0.8. This further reduced the complexity of the matrix, yielding a matrix of 871 unique domains used for training and further analysis.

### Machine Learning Approaches

To find and visualise the covariance between the domain content and pathogenicity, Partial Least Squares Discriminant Analysis (PLS-DA) was applied to the data set. PLS-DA suggested that overall, the strain specific domain content provided a good way t o discriminate between both classes, but also suggested that some strains might be misclassified. (Figure 3a & 3b).

**Fig. 3.**
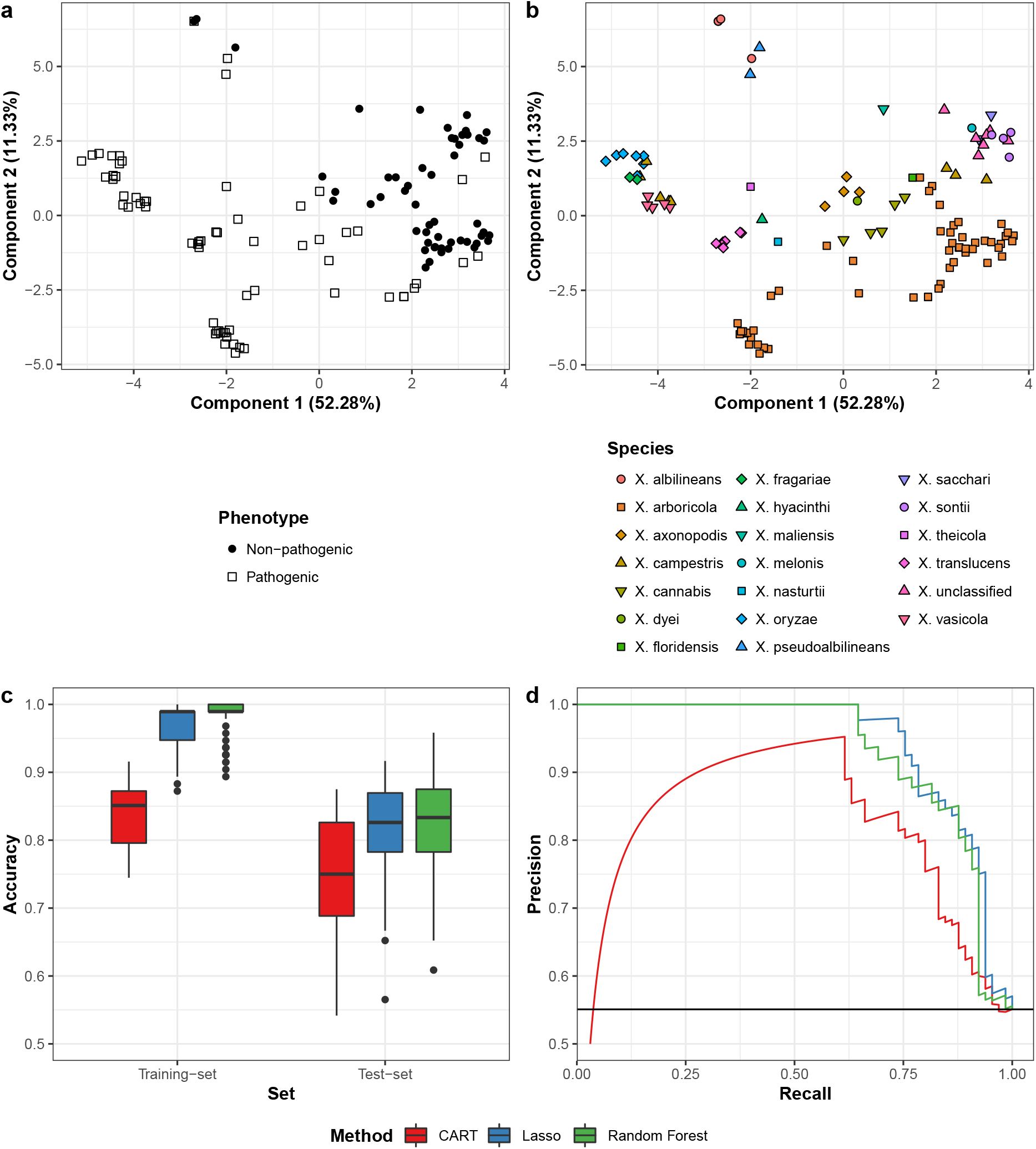
Properties of the training data set and model prediction performances. Upper panel: Partial Least Squares Discriminant Analysis (a) *Xanthomonas* strains labelled by phenotype. (b) *Xanthomonas* strains labelled according to species classification. Lower panel: Performance of the ML classifiers. (c) Classifier accuracy using a nested cross-validation scheme for each classification method. (d) Precision-Recall curves of each classification method calculated from the median test-set probabilities.

To learn more about the relationship between the domain content and the plant-associated phenotype, three other types of ML approaches, selected for their high level of interpretability and their performance on data sets with a modest number of observations, were applied: CART, Lasso, and RF.

#### Model performances

To evaluate the performance of the respective models a 20×5-4×10 nested repeated Cross Validation (CV) was used, based on recommendations of Krstajic *et al.* (57) and Kuhn *et al.* (58). To allow for a better comparison all ML approaches were trained and tested on the identical data partitions. For all approaches, the test-set accuracy was highly variable with a difference in accuracy larger than 0.3 depending on the specific data training- and test-set partition (Figure 3c). This underlines the need to estimate the variation in the model performance using nested CV, if these estimates are to be used as an indication of real world performance. The CART model showed the lowest performance with a median accuracy of 0.750. The Lasso and RF models performed similarly better, with a median accuracy of 0.826 and 0.833 respectively. The RF model performed better on the prediction of non-pathogenic strains, with an sensitivity of 0.769 and specificity of 0.909 (considering pathogens as the positive class), whereas the Lasso’s performance was more balanced, but slightly in favour of the pathogens with a sensitivity of 0.846 and specificity of 0.818. The differences between the training- and test-set performance were larger for the Lasso and RF models in comparison to the CART. The median selected tuning parameters indicated that all classifiers favoured relatively low complexity models; *cp* = 0.068 for CART (yielding 2–3 domains per tree), *λ* = 7.85 10^−3^ for Lasso (yielding ~ 29 domains with a non-zero coefficient), and *mtry* = 89 for RF (Supplementary File S3). The median precision-recall (PR) curve (Figure 3d) provided a more detailed representation of model performance. The PR-curves confirmed that the CART model underperforms in comparison to the other models. The drop in precision at low recall values of the CART is caused by the model attributing the highest probability of pathogenicity to two *X. axonopodis* strains that according to literature were non-pathogenic.

#### Species Specific Prediction Performance

To gain a better understanding of model behaviour, the *in silico* predicted class probabilities were compared with the *in vivo* predictions obtained from literature. To this end the median class-probabilities over the 20 repeats of the nested CV and the pathogenicity labels extracted from the database, were mapped onto the first two components of the previously created PLS (Figure 4).

**Fig. 4.**
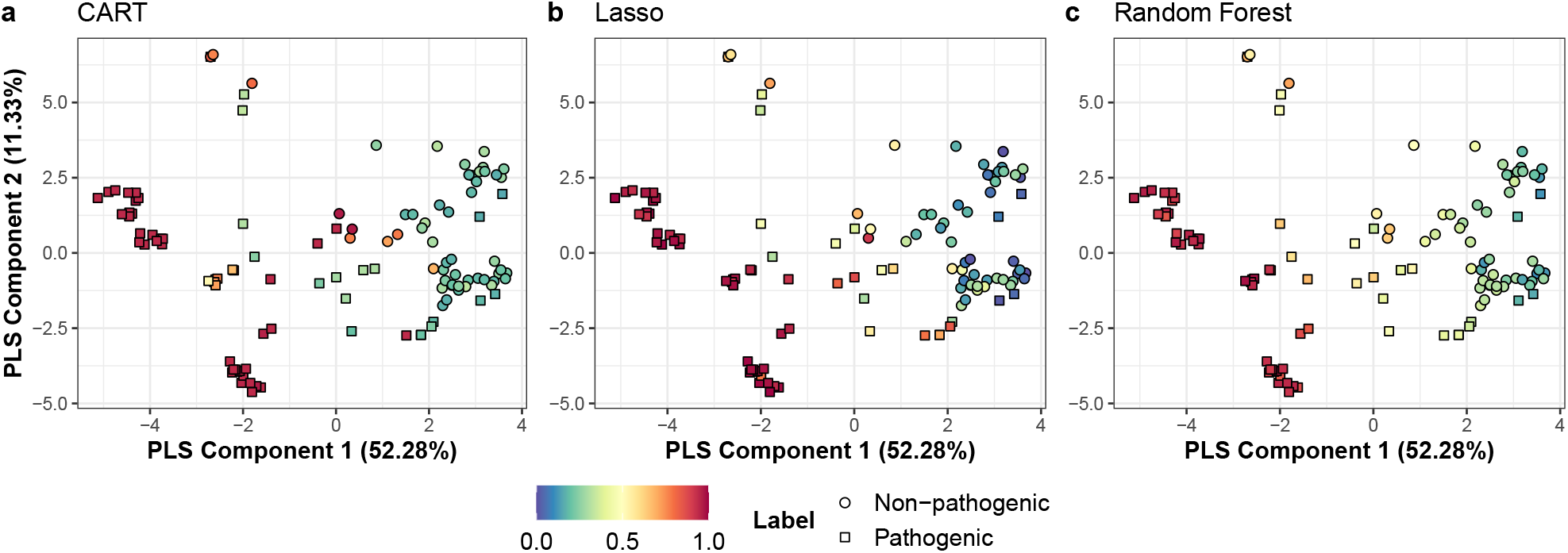
Median predicted class probabilities mapped onto the PLS-DA. Colour scale (0–1) represents the median predicted probability for pathogenicity. Labels represent the phenotype according to literature. Class probabilities were predicted for the hold out samples in the 5-fold CV outer-loop and the median was taken over the results from the 20 repeats.

Strains highlighted in red were predicted to be pathogenic with a high probability. Strains that scored consistently high across all approaches belonged to the *X. oryzae, X. vasicola* and *X. fragariae* species or to the *juglandis, corylina* and *pruni* pathovars of *X. arboricola*. For these species, with exception of *X. arboricola*, the dataset only contained genomes for strains that were pathogenic according to literature. The same was true for the *X. translucens* species, however only the Lasso and RF model predicted this species with high confidence, whereas the CART model showed mixed predictions for this species. The *X. hyacinthi, X. nasturtii* and *X. theicola* species also consisted entirely of pathogenic strains, but these species were predicted to have a lower probability of being pathogenic, which can likely be attributed to being represented with a single sequenced strain. The *X. nasturtii* strain was correctly predicted as pathogenic by all ML approaches, whereas the *X. hyacinthi* and *X. theicola* were incorrectly predicted by the median CART and Lasso models.

Strains highlighted in a blue/green colour were predicted to be non-pathogenic with a high probability. These strains formed two distinct clusters: one cluster in the lower-right containing the non-pathogenic *X. arboricola* strains and one cluster in the upper-right, containing strains from the non-pathogenic *X. sontii, X. sacchari* and *X. melonis* species and non-pathogenic strains with an undefined species taxonomy. The confidence in the predictions of the non-pathogenic strains was lower compared to the strains that were consistently predicted as pathogenic, as indicated by a weaker deviation from the neutral score of 0.5. This increased uncertainty likely stems from four strains that are pathogenic according to literature, but are located close to the clusters of non-pathogens.

Strains located near the origin and top-middle of the PLS mainly consisted of pathogenic strains from the remaining *X. arboricola* pathovars and of the species with a small number of genomes with a mixed phenotype (*X. cannabis, X. axonopodis* and *X. (pseudo) albilineans*). All ML approaches were uncertain about the *X. albilineans* species located at the top of the PLS, as indicated by the neutral probabilities. The species near the origin of the PLS showed a large difference between approaches. The CART model failed to correctly predict strains from both the *X. cannabis* and *X. axonopodis* species, the Lasso performed better on the *X. cannabis* species whilst still failing to reliably predict the *X. axonopodis* species and the RF gave neutral predictions for both species. The CART model also performed poorly on the less successful pathovars of the *X. arboricola* species. The Lasso slightly outperformed the RF on these *X. arboricola* pathovars, although the predictions made by the Lasso were more variable between strains.

### Feature importance

All three ML approaches apply a form of feature selection, yielding variable importance scores for features and providing insights in the classifier’s decision making process. For each ML approach these scores were obtained in a different way. For CART, variable importance was obtained by tabulating reduction in loss function for all candidate variables considered at each split; for Lasso, variable importance was computed from coefficients using t-statistic; for RF the mean decrease in accuracy when a given variable was left out of bag was used as a measure of variable importance. Scores were summed over all folds of the repeated CV outer-loop and were scaled to have the most important domain at 100. For comparison purposes, domain enrichment was calculated for each class using a two-side Fisher exacted test with Benjamini-Hochberg correction. The top 10 most important domains for each of the ML approaches were combined and sorted by enrichment score (Table 4). To provide additional context, the table also includes highly correlated domains removed by matrix filtering. The top 10 domains of CART and RF show a strong correlation, whereas Lasso used more less enriched domains.

**Table 4.**
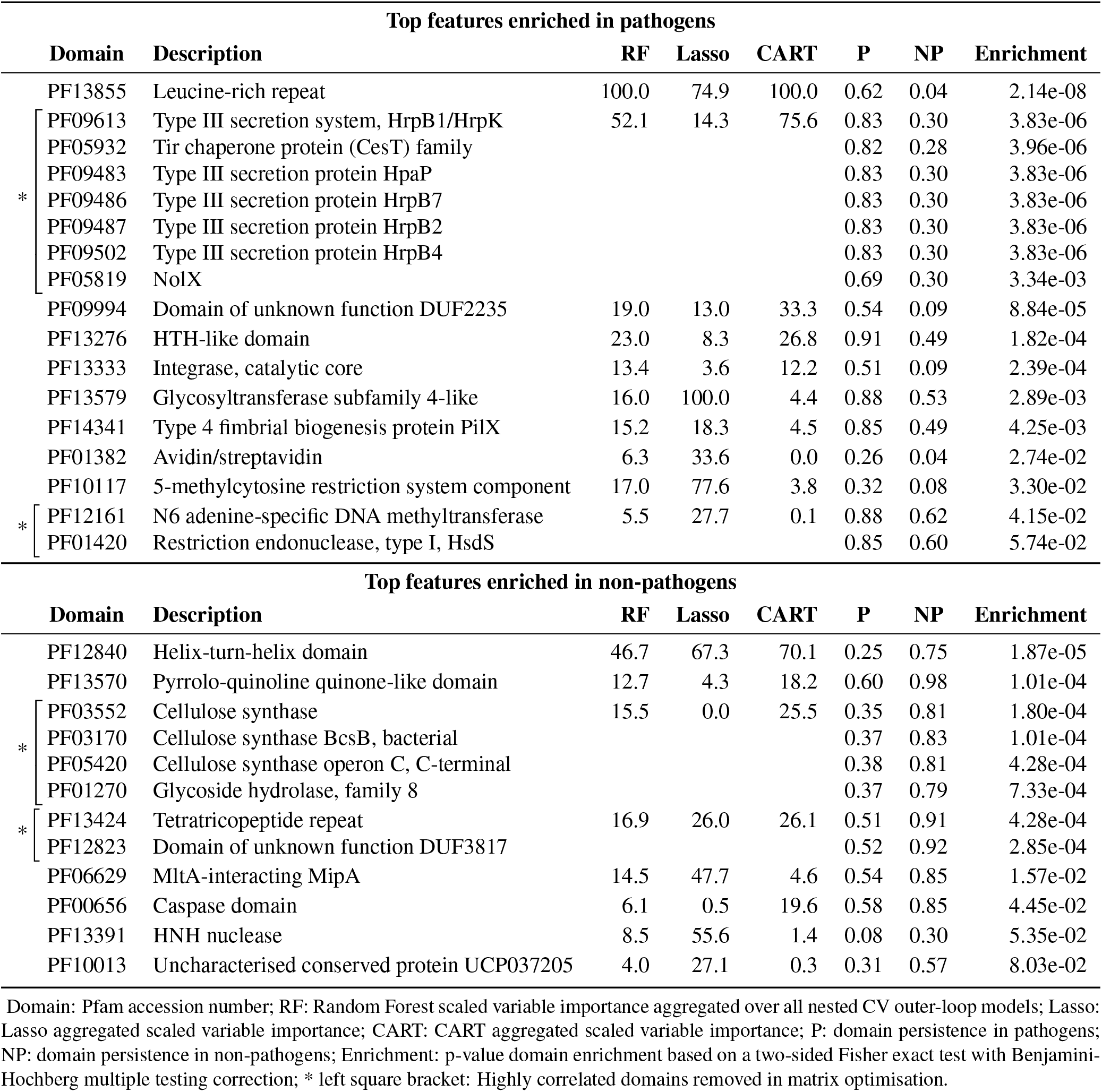
Top features used in the classifiers.

## Discussion

The *Xanthomonas* genus encompasses a diverse set of species able to infect a large variety of important crops. As non-pathogenic strains seems to constitute a significant part of the *Xanthomonas* population, from a pest-control point of view the need arises to develop means to reliably distinguish this phenotype, while the genomic diversity may shed light to the *Xanthomonas* plant-associated lifestyles and contributing traits. Currently more than 1700 *Xanthomonas* genomes are available in public repositories, which enables the application of computational approaches for studying the genomic basis for pathogenicity. However, such approaches are hampered by the current publication bias towards pathogenic strains and fragmentation of information regarding the plant associated phenotypes of *Xanthomonas* specific strains. In order to obtain a reliable and balanced set of meta-data representing both plant-associated phenotypes, we collected pathogenicity assays from recent studies that included both plant-associated phenotypes in their study and stored this information in a relational database. Strains unable to induce symptoms on all tested hosts, including the isolation host, were considered to be non-pathogenic. Still, we expected that some mislabelling of the training data was inevitable: First, because the relative abundance of the non-pathogens in nature is unknown and second, due to a high dependency of host susceptibility on the *in vitro* conditions, while there is no complementary test for non-pathogenicity.

Available Genome sequences were retrieved from GenBank and *de novo* annotated for Pfam domains using the SAPP platform (54) A heap analysis of the resulting domain matrix estimated that the pan-domainome was closed, indicating that the data set contained the majority of the variable domains present in the *Xanthomonas* genus and despite some evidence for mislabelling, a partial least squares discriminant analysis indicated that a good discrimination between both phenotypes is possible when a diverse set of domains is considered. To enable inclusion of multiple domains into the decision making process, three different machine learning approaches were explored.CART and Lasso favour the lower-complexity models with the median models using 3 and 29 protein domains respectively. By design, RF used nearly all domains, but the median tuning parameter *m_try_* = 89 was higher than default 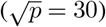 indicating that there are only a limited number of important features.

Overall, non-pathogens were classified with a lower level of certainty by all ML approaches. A large part of this uncertainty seemed to stem from four pathogenic strains that showed a strong similarity to many of the non-pathogens according to the PLS-DA. Upon closer examination, the identification of these four strains as pathogens seemed doubtful: *X. sontii* strain ASD011, was the only pathogenic member of a species that has previously been defined by it’s non-pathogenicity (5). *X. campestris* NCPPB4393, actually belonged to the *X. sacchari* species, a species of which the pathogenicity is still ill defined (59) and this strain in particular was special, as it is the only *Xanthomonas* strain known to be isolated from an insect host. *X. arboricola* LMG19145 belonged to the *X. arboricola* pv. *fragariae* subspecies, a subspecies with many conflicting pathogenicity reports, that have not been resolved to date (13, 14). *X. arboricola* 3004 was an aberrant strain that didn’t cluster with any of the pathovars in the *X. arboricola* species and this strain was only known to be weakly pathogenic to barley. A closely related strain, *X. arboricola* CITA 44, was not able to cause any symptoms on this same host or any other tested host (29). This indicates that these strains are either mislabelled in literature, or belong to a class of very weak opportunistic pathogens. There also remains the possibility that strains identified as non-pathogens by literature symbiotically rely on co-infection with pathogenic species (15). Given that infection assays typically use pure cultures, such lifestyle would go unnoticed.

Almost half of the tested strains belong to the *X. arboricola* species (Table 2). The predicted class probabilities correlated strongly with varying levels of pathogenicity reported in literature (10). Within this species the strains belonging to the highly pathogenic pathovars (*juglandis, corylina* and *pruni* pathovars) or non-pathogenic strains were correctly identified with a high probability. Strains belonging to the weakly pathogenic pathovars obtained more neutral probabilities (Figure 4) which might suggest that, at least for *X. arboricola*, these models can not only combine multiple features to discern phenotypes but are also able to score different levels of pathogenicity.

### Important features enriched in the pathogens

Extraction and examination of the most important features used for prediction by the respective ML approaches, showed that the CART and RF considered mostly the same domains, with both favouring domains that were highly enriched in one of the classes. The Lasso behaved distinctly different with its variable importance scores showing a weaker correlation with the enrichment analysis.

Many of the highly important domains that were enriched in pathogens have already been related to pathogenicity in literature, indicating that the here used approach is capable of detecting biologically relevant features. Overall, the most important feature across models was a leucine-rich repeat (LRR) domain present in two *Xanthomonas* type III effector proteins (XopL and XopAE/HpaF). The domain was present in nearly two-thirds of the pathogens and absent in the non-pathogenic strains, with exception of two potentially mislabelled *X. axonopodis* strains. Similarly, a large group of correlated domains representing the Type III Secretion System (T3SS) was also found to be highly important, although these domains had a much broader representation amongst both phenotypes. The T3SS and its effectors are is the most important and extensively studied aspect of *Xanthomonas* pathogenicity (16). The entire T3SS is encoded by the *hrp* cluster and consists of more than 20 proteins that form a needle-like syringe used to inject proteins into the host cytoplasm (60). Type III effector proteins are translocated into host cells where they target many host components, serving to suppress the host immune system, increase nutrient availability and facilitate the infection process (17). Specifically, mutations in HpaF were found to impact virulence in *X. axonopodis* (61).

Amongst the important predictors enriched in pathogens were two domains related to mobile genetic elements. The first, P F13276, was a helix-turn-helix-like domain t hat was found in several IS3 transposases and the second, PF13333, was an integrase domain that belonged to a putative OrfB transposase. Whilst both domins were enriched in pathogens, the the helix-turn-helix-like domain was also well represented in the non-pathogens, whereas the integrase domain was only found in a few non-pathogens and was more scattered across the pathogens. Transposases are known to flank pathogenicity islands in genomes of *Xanthomonas* (62). However, these domains are more commonly found within pathogenic islands, where, in some cases, they are assumed to have played a role in the initial mobilisation or subsequent rearrangement of the element (63).

The most important domain for the Lasso model was PF13579, a n-terminal glycosyl transferase 4-like domain of the RfaB family. This domain is most likely involved in LPS production, which is important for pathogenicity by providing a barrier against anti-microbial compounds, facilitating adhesion and preventing host recognition (15). Although the domain was enriched in pathogens, it was found pathogens and non-pathogens of all major species, with the exception of *X. albilineans*.

Next to domains that were already known to be involved in pathogenicity, a domain of unknown function (DUF2235), enriched in pathogens, was also found to be an important predictor. The domain, which represents a further uncharacterized alpha/beta hydrolase, was present in up to 16 proteins per genome (7 on average). Further analysis revealed that in one cluster this domain was fused with domain PF16014 (histone deacetylase complex subunit SAP130 C-terminus domain). This domain is usually found to be part of transcriptional repressor Sin3 and the region containing this domain was also flagged as a superantigen-like protein SSL3 motif. The SSL3 protein is important for pathogenicity in the human pathogen *S. aureus*, where it is known to bind to the hosts Toll-Like Receptor 2 (TLR2), inhibiting stimulation by its ligands (64) suggesting an important role for suppressing the plant hosts immune system.

### Important features enriched in the non-pathogens

Domain enriched in non-pathogens or notably absent from pathogens were also highly important for model predictions. Many of these domains have an implied role in increasing resistance against environmental factors, which is in line with the idea that non-pathogenic strains are generalists that can survive in a much broader range of conditions than their pathogenic counterparts (11, 12). All methods agree that another Helix-Turn-Helix domain (PF12840) is an the important discriminant enriched in non-pathogens. This domain was found in all non-pathogens with exception for a subgroup of *X. arboricola* and in some pathogens of the *X. oryzae* and *X. campestris* species. The domain is found in DNA binding transcriptional regulators of the ArsR/SmtB, Arsenical Resistance Operon Repressor, family. In *X. campestris* 8004 it was found that the HTH ArsR containing gene is upstream of arsenite efflux pump AcR3 and a putative high-affinity *Fe*^2+^*/Pb*^2+^ permease. In the same strain it was shown, via a knockout, that the *arsR* gene confers an increased resistance against arsenate (65).

Amongst the important predictors enriched in non-pathogens, was a group of correlated domains involved in cellulose synthesis. The model organism for studying bacterial cellulose synthase is *Acetobacter xylinum*. For this organism it is known that the complex produces and transports beta-1,4-glucan chains, creating rigid crystalline structures on the outer membrane. Bacterial cellulose can fulfil diverse roles from mechanical/environmental protection to cell adhesion during symbiotic or pathogenic interactions (66). Cellulose synthesis is promoted by cyclic-di-GMP trough the PilZ domain present in glycosyltransferase CeSA, which is part of the membrane-bound cellulose synthase complex (67). Cyclic di-GMP is also known to down-regulate biofilm formation, EPS production, extracellular enzyme production and *hrp* gene expression in *Xanthomonas* (15). Thus given that cellulose synthesis and production of pathogenicity factors, is inversely related we hypothesise that these domains might be involved in providing environmental protection when the bacterium is not shielded by the host homeostasis. The last domain that could be linked to environmental resistance was the MltA-interacting MipA domain (PF06629) was found in all *Xanthomonas* species, but had a lower presence in the highly pathogenic *X. arboricola* pathovars. MipA is a protein that mediates assembly of MltA into the PBP1B murin transglycosylase/transpeptidase complex. Mutations in other genes of the *mlt* family are related with morphological abnormalities in *X. campestris* (68). Given that the domain is widely present in both pathogenic and non-pathogenic strains, it seems unlikely that the domain is central to non-pathogenicity. However, the morphological changes induced by loss of genes containing this domain, could impair the bacterium’s ability to resist mechanical stress.

## Conclusion

The plant-associated phenotype of a *Xanthomonas* strain is the result of a multitude of non-persistent traits. Consequently a single genome encoded feature will have limited power to predict the plant-associated phenotype. By using machine learning methods that take into account an ensemble of domains, a better prediction can be obtained. Through recursive feature elimination, key domains related to plant-associated phenotype could be identified leading to the discovery of novel traits.

## Methods

### Data processing

Data processing, analysis and model building was done in *R* (v4.0.5). SQL was used to communicate with the phenotype database and SPARQL was used to communicate with the Graph annotation database. All R scripts were executed on a Windows 10 machine.

### Phenotype Database

Pathogenicity assays of individual *Xanthomonas* strains were mined from literature that also considered non-pathogenicity. Relevant parameters and outcomes were stored in a relational database MySQL database (*MariaDB* v10.5.9). To create the database, the database model was forward-fengineered into the *create_database.sql* script using *MySQL Workbench* (v8.0.24). A manually curated Excel form (*Supplementary File S1*) was used o fill the database and subsequently loaded into the database using the *input_data.R* script. Connection to the database was established using the base R *DBI* library and the *RMariaDB* (v1.1.0) driver. The script was also used to enforce additional constraints on the values of specific fields.

### Data Retrieval

The *genbank.R* script was used to interface with the phenotype database. For the strains with a known pathogenicity, genomes were retrieved directly from the *NCBI Genbank* genome repository (accessed May 6th, 2020) using *RCurl* (v1.98-1.2). If multiple genomes were available for a single strain, the genome with the highest quality of assembly was taken.

### Genome Annotation

Genomes were de novo annotated using the *SAPP* framework (54), running on a Linux machine (openSUSE leap 15.1) with *OpenJDK* v11.0.5. The retrieved genomes were converted to a HDT format using the *Conversion* module from SAPP (69). Protein encoding genes were identified using Prodigal (55) and annotated for Pfam protein domains (70) using *InterproScan* (v5.44-79.0) (56). Fully annotated genomes and their provenance, were uploaded to a linked data repository using *GraphDB* (v9.7.0) for further analysis. To speed up computation SAPP modules were run in parallel using the *GNU Parallel* CLI (71).

### Data Analysis and Visualization

Annotation results were retrieved from the linked data repository using SPARQL queries and the *SPARQL* R package (v1.16). The binary Pfam domain presence/absence matrix was generated with (*sparql.R*). Dendrogram: Distances were calculated using the base R *dist* function with a Manhattan distance measure and hierarchical clustering was performed using the base R *hclust* function with average linkage. The *ape* (v5.4-1) R package was used to root the tree. The resulting dendrogram was visualised using the *dendextend* package (v1.14.0). The enrichment of single domains between and the two phenotypes was tested using a two-sided Fisher exact test with Benjamini-Hochberg multiple testing correction, using the R base *fisher.test* and *p.adjust*.

### Heap analysis

The *micropan* (v2.1) package (72) was used. The effect of sample size on the estimated/observed pan and core domainome sizes was explored by repeatedly (*n* = 100) sampling a fixed number of genomes using 20 different sample sizes equally distributed over the range spanning from 5 to 118, with 118 being the total number of genomes in this stduy. For each sample, the observed pan and core sizes were inferred directly from the data and the estimated pan and core sizes were obtained by fitting a binomial mixture model (with *k* ranging from *k* = 3 to *k* = 17) to the selected subset and taking the estimate with the lowest BIC.

### Matrix optimization

Domains with near-zero variance were removed using a threshold of present or absent in >97.5 genomes. For highly correlated domains a representing domain was chosen based on a Pearson correlation of *ρ* > 0.8.

### Partial Least Squares Discriminant Analysis

A Partial Least Squares (PLS) Discriminant Analysis was performed by training a two-component PLS regression model on domain matrix using the “pls” (v2.7-3) R package. The first two-components were then extracted and visualised.

### Model development

Model tuning and testing (*model_building.R*) was performed using the *Resample-Model* function, contained in custom a R library created for this analysis. This library provided an interface between the *rsample* package (v0.0.9), used for data partitioning, and the *Caret* package (v6.0-86), used for model building and tuning (73). The filtered pathogenicity dataset was partitioned using a nested Cross-Validation (CV) scheme consisting of a 20-times repeated 5-fold outer-loop and a 4-times repeated 10-fold inner-loop. Accuracy was used as the performance metric in both loops and additional statistics were generated using Caret’s *twoClassSummary* function. When calculating calculating binary classification metrics pathogenicity was regarded as a ‘positive’ result and non-pathogenicity as a ‘negative’ result.

Using identical partitions, different types of models were built and tested: CART models from the *rpart* package (74), Lasso models from the *glmnet* package (by setting *α* = 1) (75), and Random Forest models from the *RandomForest* package (76). For the CART models, the complexity parameter *cp* was varied across Caret’s default grid, with a size of 9. For the lasso model the parameter *λ* was optimised using a grid ranging from 1 10*−*4 to 1 with a size of 20 and an exponentially increasing step size. For the random forest model, the number of trees was fixed at 1000 and *mtry* was varied over Caret’s default grid of size 9.

### Model Performance & Variable Importance

The performance of the respective approaches on the different genomes was examined by superimposing the median probability for a strain to be pathogenic over the previously created PLS-DA visualisation. For each approach, the variable importance scores for per domain were calculated using Caret’s build-in *varImp* function with scaling turned off. Results from multiple folds were combined by summing the unscaled variable importances for each domain, after which the final results were scaled to have the most important variable at 100.

## Supporting information

The code and SQL database are available at: https://gitlab.com/wurssb/xanthomonas-phenotype-prediction.

Genome annotations in a binary RDF format (HDT) is available at http://doi.org/10.4121/14546625.

**Supplementary Figure S1 (PDF)**: Entity-Relationship diagram of the *Xanthomonas* phenotype database.

**Supplementary Figure S2 (PDF):** Annotation statistics of 118 sequenced *Xanthomonas* strains used in this study.

**Supplementary Figure S3 (PDF):** Domain based distance tree of the 118 *Xanthomonas* strains used in this study.

**Supplementary File S1 (XLSX):** Input and output of the SQL database.

**Supplementary File S2 (XLSX):** Genome metadata, annotation statistics and the domain matrix.

**Supplementary File S3 (XLSX):** Model statistics and domain enrichment.

## Author contributions

D.M., W.P., P.S., J.K. participated in the conception and design of the study. D.M. was responsible for the code and design of the database. D.M., W.P., P.S., J.K. wrote sections of the manuscript. All authors critically revised the manuscript.

---

**Fig. S1.**
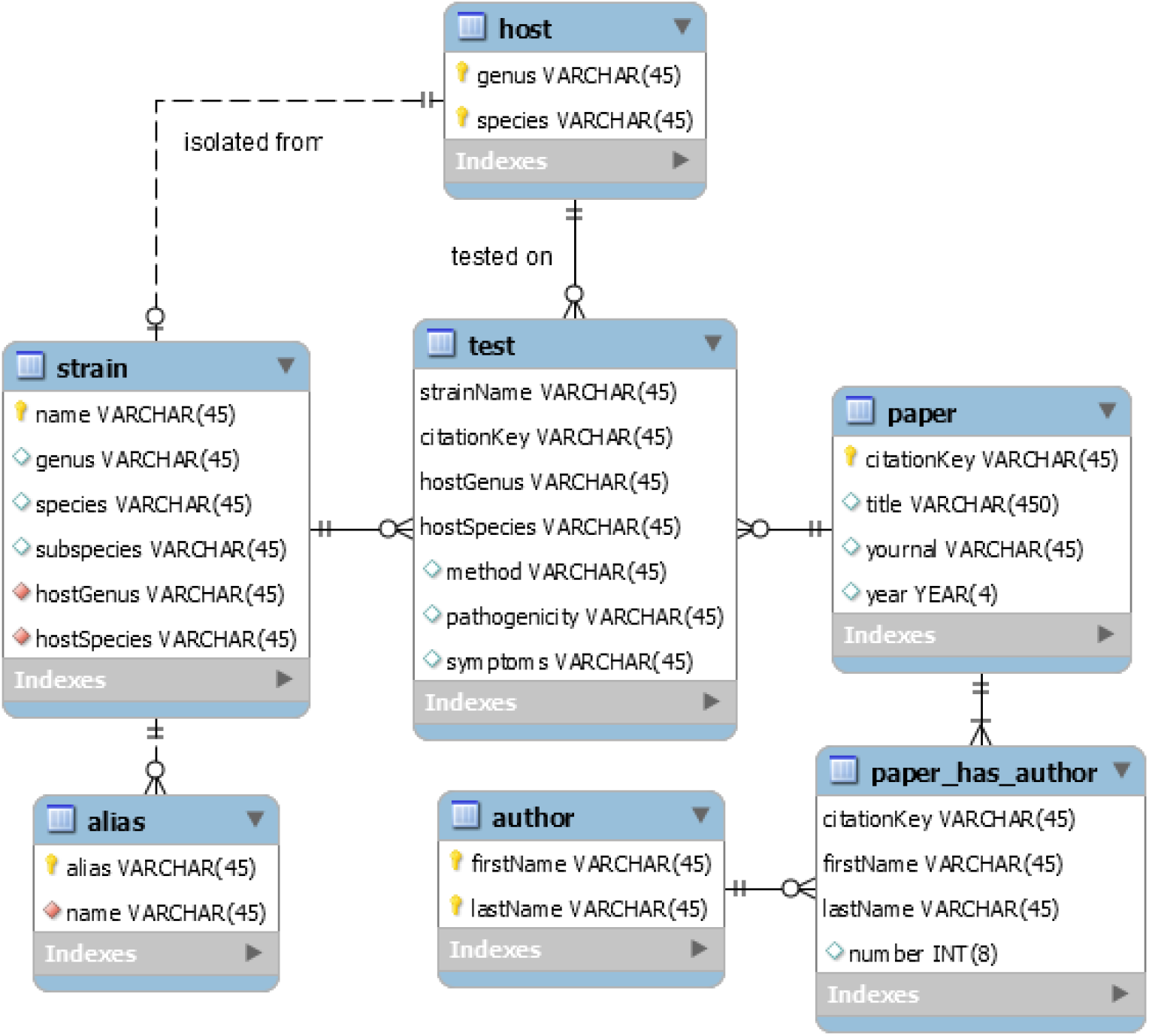
Entity-Relationship diagram of the *Xanthomonas* phenotype database. Solid line: dependent relationship, dotted line: independent relationship.

**Fig. S2.**
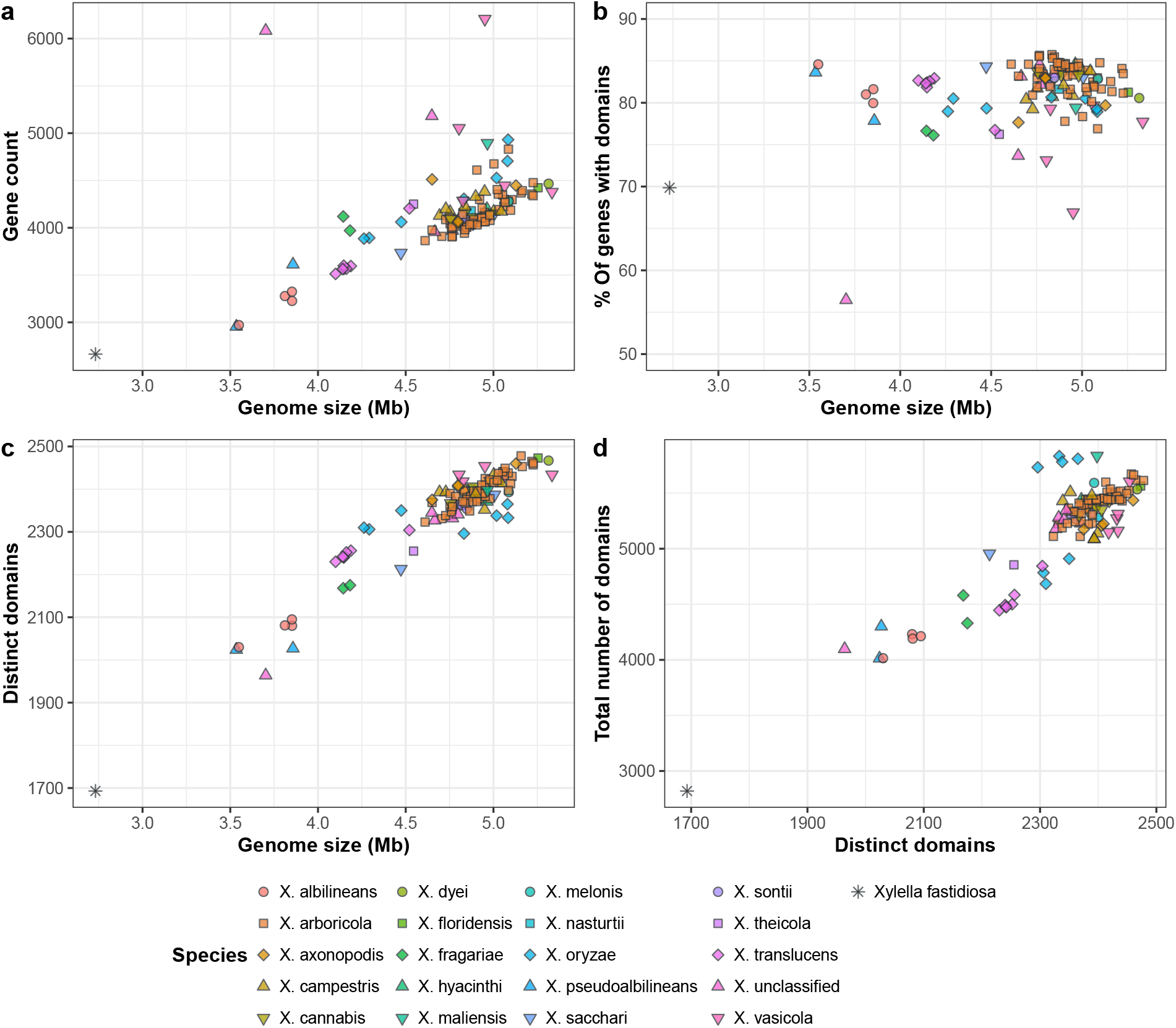
Annotation statistics of 118 sequenced *Xanthomonas* strains used in this study. Xylella fastidiosa 9A5C (marked by *) was used as an out-group.

**Fig. S3.**
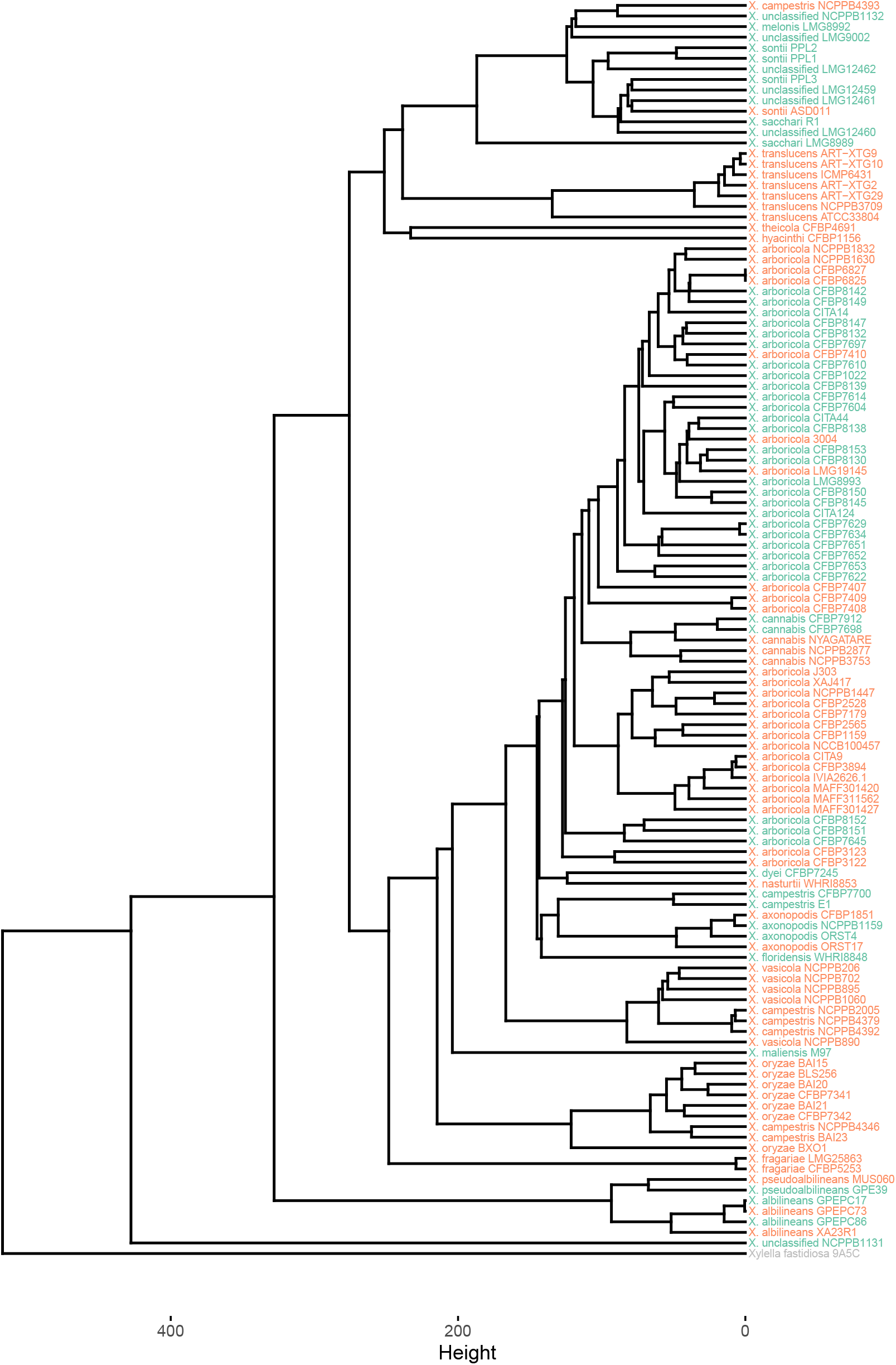
Domain based distance tree of the 118 *Xanthomonas* strains used in this study. The tree was calculated on the binary domain presence/absence matrix using Manhattan distance. Colours indicate pathogenicity according to literature: red = pathogenic; green = non-pathogenic; Xylella fastidiosa is used as out-group (in grey).

